# Coral Guard substrates accelerate growth and fragment fusion for coral restoration

**DOI:** 10.64898/2026.07.21.739923

**Authors:** Stefan Kolle, Netanel Kramer, Christopher Suchocki, Joshua L. Kuailani, Lindsey Badder, Natalie Levy, Sofia P.C. Martin, Aiden Beaupain, Alexander Valle Perez, Samapti Kundu, Eric Schuster, Christopher B. Wall, Rob Toonen, R3D consortium, Daniel Wangpraseurt

## Abstract

Coral reefs are declining globally, creating an urgent need for scalable technologies that improve the efficiency and effectiveness of active reef restoration. Coral reef restoration is increasingly constrained by algal overgrowth, which suppresses coral growth, survival, and restoration efficiency. Building on the recently developed Coral Guard platform, we evaluate its restoration performance under long-term *in situ* coral nursery conditions and introduce Fusion Guard Tiles, a geometry-optimized Coral Guard design that accelerates microfragment fusion. We evaluated Coral Guard Plugs using the branching coral *Stylophora pistillata* under complementary *ex situ* conditions and Fusion Guard Tiles using the massive reef-building coral *Porites evermanni* in *in situ* coral nurseries. In *P. evermanni*, Fusion Guard Tiles increased lateral tissue growth 2.6-fold and three-dimensional tissue surface area growth by >2.7-fold after 6 months compared with conventional substrates. After 12 months, colony height and volume were approximately 4.0-fold and 2.2-fold greater, respectively, while complete fragment fusion occurred only on Fusion Guard Tiles. In *S. pistillata*, Coral Guard Plugs increased lateral tissue growth by ∼1.7-fold. Microcomputed tomography revealed 10–13% higher skeletal density in both species. In *P. evermanni*, Fusion Guard Tiles also increased symbiont density by 70% and tissue protein content by ∼2.5-fold relative to controls. Together, these findings demonstrate that Coral Guard substrates suppress algal competition while accelerating coral growth, skeletal development, and microfragment fusion, providing a scalable, low-maintenance technology to enhance coral nursery productivity and reef restoration.

## Introduction

Coral reefs are among the most biologically diverse and socioeconomically important ecosystems on Earth, supporting approximately 30% of marine biodiversity and sustaining the livelihoods of nearly one billion people while occupying less than 1% of the ocean surface (Duarte et al. 2025; Fezzi et al. 2023; Hughes et al. 2002). In addition to supporting fisheries and tourism economies, coral reefs provide critical coastal protection by dissipating wave energy and buffering shorelines from storm impacts (Ferrario et al. 2014). Yet, well over half of global live coral cover may already have been lost, and projections indicate that even at 1.5°C warming, reefs could decline by a further 70 to 90 percent, with losses exceeding 99% at 2°C (Bellwood et al. 2004; Duarte et al. 2025; Eddy et al. 2021). Recurrent marine heatwaves, ocean acidification, and deoxygenation interact with local stressors such as eutrophication, pollution, and overfishing to suppress recovery and amplify mortality (Alderdice et al. 2022; Pezner et al. 2023; Zaneveld et al. 2016). Even under ambitious mitigation scenarios, substantial and potentially irreversible coral loss is expected due to committed warming and ecological lag effects (Duarte et al. 2025; Lenton et al. 2025).

In this context, active intervention is required to sustain the structure and function of reefs. Coral reef restoration encompasses approaches such as transplantation, nursery propagation, and larval enhancement (Boström-Einarsson et al. 2020; Peixoto et al. 2024, Segaran et al. 2024). While these interventions can increase coral cover when key bottlenecks are addressed (Lange et al. 2024), many restoration efforts remain constrained by low throughput, high labor demand, and limited scalability (Boström-Einarsson et al. 2020). A major bottleneck in fragmentation-based restoration is the slow growth of coral fragments, driven in part by intense competition at the coral– substrate interface (Karimi et al. 2025, Lustic et al. 2020). Benthic algae rapidly colonize available substrate, directly competing with juvenile coral recruits and adult outplants for space and light (Van Woesik et al. 2018). In addition, benthic algae also alter local biogeochemical and microbial conditions through increased dissolved organic carbon release and microbial activity (Barott & Rohwer 2012; Haas et al. 2011; Haas et al. 2016). These interactions suppress lateral tissue expansion and delay fragment fusion, limiting the formation of larger, structurally stable colonies. Improving early growth dynamics in coral outplants represents a key opportunity to enhance restoration efficiency, as accelerating lateral tissue expansion and promoting earlier fragment fusion can reduce nursery residence time, increase post-outplant survivorship, and improve scalability (Page et al. 2018; Knapp et al. 2022; Schmidt-Roach et al. 2025). Critically, current coral gardening practices rely on frequent, labor-intensive cleaning to maintain fragment performance, making maintenance a primary bottleneck to scale (Schmidt-Roach et al. 2025). Reducing fouling while minimizing maintenance therefore represents a major opportunity to increase restoration efficiency and scalability. Despite advances in coral genotype selection, husbandry, and nursery design, the substrate microenvironment that governs early growth remains comparatively underexplored. Directly engineering this interface offers a clear pathway to scalable restoration by targeting underlying ecological processes in a manner that can be robustly deployed across variable environmental conditions (Baer et al. 2025).

Applying bioengineering principles enables intentional modulation of the microenvironment surrounding coral outplants to shift competitive and physiological dynamics in favor of coral growth (Valenzuela Matus et al. 2024; Jia et al. 2024; Karimi et al. 2025; Corigliano et al. 2025). Recently, the Coral Guard platform was developed as a fouling-release restoration substrate that suppresses algal competition while allowing coral tissue to overgrow the material, demonstrating proof-of-concept under laboratory conditions (Karimi et al. 2025). However, whether Coral Guards improve coral growth and restoration performance under long-term *in situ* coral nursery conditions remains unknown. Here, we evaluate the restoration performance of the Coral Guard platform through complementary *ex situ* and long-term *in situ* coral nursery experiments. We further introduce Fusion Guard Tiles, a geometry-optimized Coral Guard design developed to promote rapid microfragment fusion in massive corals. Using the branching coral *Stylophora pistillata* and three key Hawaiian restoration species (*Porites evermanni*, *Montipora capitata*, and *Pavona varians*), we evaluate colony growth, fragment fusion, skeletal development, tissue biomass, and coral physiology through complementary *ex situ* experiments and long-term *in situ* coral nursery deployments.

## Methods

### Coral Guard Platform Fabrication and Fusion Guard Tile Design

For this study, Coral Guard substrates were fabricated following the previously described protocol (Karimi et al. 2025). Briefly, PDMS (polydimethylsiloxane) coatings were prepared using a SYLGARD™ 184 Silicone Elastomer Kit at a 10:1 elastomer base-to-curing agent ratio. Components were mixed using a ThinkyMixer ARE-310 at 2000 rpm for 60 s, followed by degassing at 2200 rpm for 60 s.

The Coral Guard platform comprised three substrate configurations tailored to different experimental applications. First, Coral Guard Plugs (3 cm diameter) consisted of fully coated aragonite-based frag plugs and were used for *Stylophora pistillata* growth experiments. Second, split-choice Coral Guard Plugs were fabricated by milling one half of aragonite-based frag plugs (Ocean Wonders, Agrocrete, item: 804879061113) to a depth of 1 mm using a CNC machine (Carbide3D Nomad 883). Only the recessed half was coated with PDMS, resulting in adjacent Coral Guard and control surfaces of equal height for within-colony comparisons (Fig. 1A). Third, for long-term *in situ* nursery experiments, we developed Fusion Guard Tiles, a geometry-optimized Coral Guard design fabricated using 4-inch Reefing Art Breakable Coral Frag Tiles. Fusion Guard Tiles incorporated four recessed attachment sites designed to hold individual microfragments while promoting rapid tissue fusion and maintaining Coral Guard surfaces between fragments. Control substrates consisted of uncoated aragonite frag plugs (Ocean Wonders) for Coral Guard Plug experiments and uncoated frag tiles of identical geometry for Fusion Guard Tile experiments.

**Figure 1.**
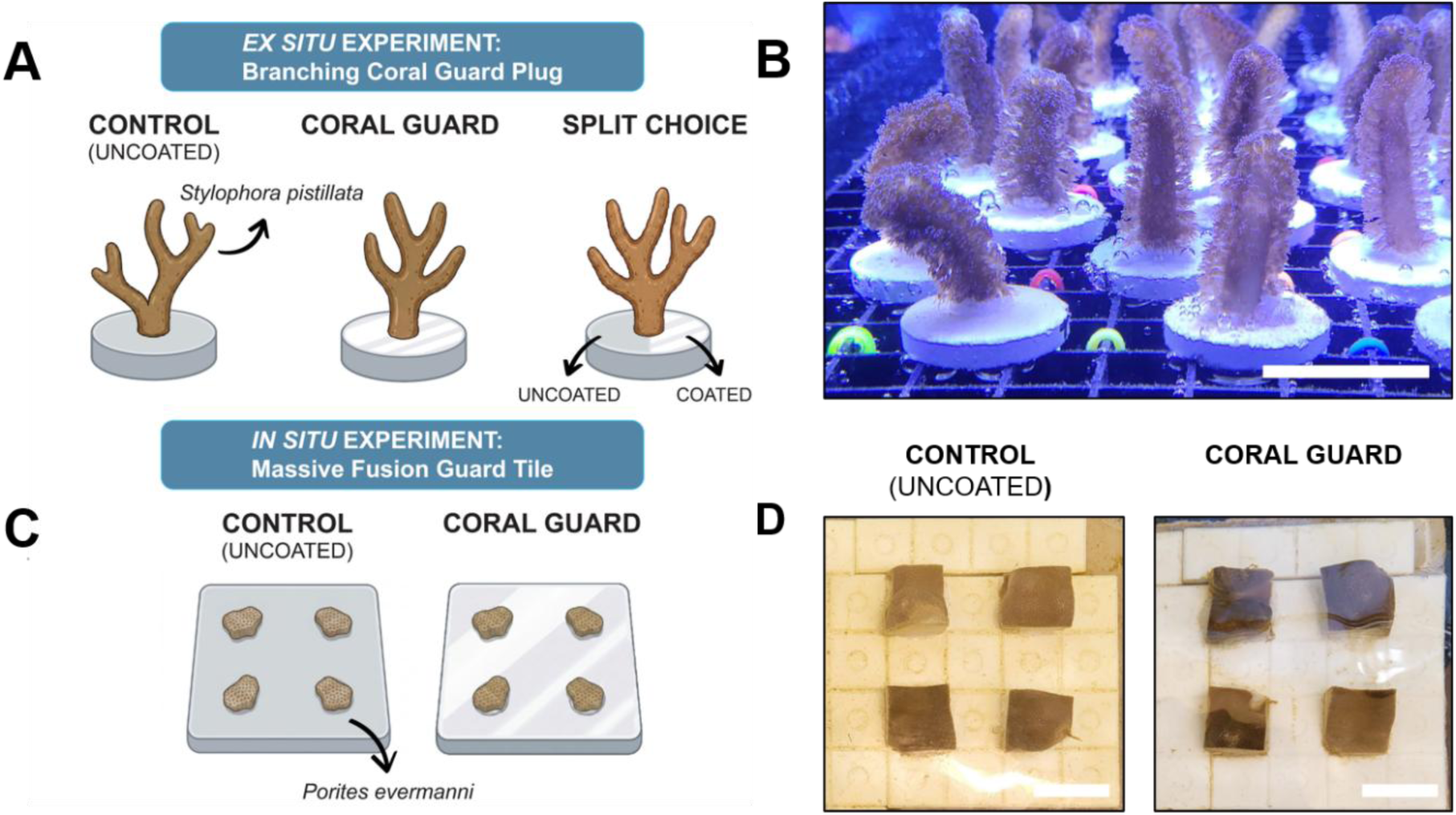
Experimental design for complementary *ex situ* and *in situ* coral growth studies. (A) Coral Guard Plug configurations used in the *ex situ* experiments, showing Control, Coral Guard Plug, and Split-Choice Coral Guard Plug treatments. (B) *Stylophora pistillata* fragments attached to the different plug treatments (scale bar = 3 cm). (C) Fusion Guard Tile design used for the *in situ* coral nursery experiments, showing four *Porites evermanni* microfragments positioned within the tile to promote fragment fusion. (D) Control and Fusion Guard Tiles following microfragment attachment (scale bar = 3 cm).

### Ex situ experimental design

*Stylophora pistillata* colonies were obtained from Birch Aquarium at Scripps Institution of Oceanography (San Diego, USA), fragmented into ∼2.5 cm pieces, and were maintained in a flow- through aquarium system at 25 °C under a controlled light-dark cycle (Fig. 1A-C). Experiments were conducted in an aquarium system supplied with natural seawater, which provided a mixed community of temperate filamentous algae. Although these algae are not common competitors in tropical reef systems, they provided a consistent model to evaluate algal fouling and coral tissue overgrowth. Coral Guard Plugs were compared with uncoated aragonite-based frag plug controls. For each replicate, a single coral fragment was attached to the center of the plug using cyanoacrylate-based aquarium glue (RA Aqua Tech, USA) (Fig. 1B). Two complementary experiments were performed. In the growth experiment, coral fragments were mounted on either Coral Guard Plugs or uncoated control plugs (n = 8 per treatment) to quantify differences in lateral tissue expansion. In the split-choice experiment, Coral Guard Plugs contained adjacent coated and uncoated surfaces, allowing individual coral fragments to simultaneously interact with both substrate types (n = 11). This design enabled within-colony comparisons of tissue growth allocation and overgrowth behavior in response to substrate type.

### In situ coral nursery experimental design

*In situ* experiments were conducted at the Hawai‘i Institute of Marine Biology (HIMB) coral nursery on Moku o Lo‘e in Kāne‘ohe Bay, O‘ahu, HI (Fig. 1D-F). Fusion Guard Tiles (4-inch square) and uncoated control tiles were mounted onto panel arrays (9 × 10 inch) (Fig. 1D), with two panels per treatment. Each tile housed four microfragments of a single species and genotype to enable controlled fusion. Three restoration-relevant coral species *Porites evermanni, Pavona varians,* and *Montipora capitata* were tested, with eight replicate tiles per species and treatment (Fig. 1E). Panels were deployed on floating nursery platforms and maintained under ambient field conditions (Fig. 1F). Coral performance was assessed at 3, 6, and 12 months using three-dimensional photogrammetry. Due to fragment detachment and coral bleaching during the study period, datasets for *P. varians* and *M. capitata* were incomplete and are therefore only reported in the supplementary information (Fig. S3, Fig. S4).

### Photogrammetry

For the *in situ* experiments, three-dimensional (3D) models of coral colonies were generated using structure-from-motion photogrammetry following established coral reef reconstruction protocols (Ferrari et al., 2017). At each sampling time point, colonies were imaged *in situ* by SCUBA using a Sony α7 III (ILCE-7M3) camera in an underwater housing. Images were acquired under ambient lighting with 50–80% overlap between successive photographs to enable high-fidelity 3D reconstruction. Image processing was performed in Agisoft Metashape Professional (v2.3.0, Agisoft LLC). Images with quality scores <0.5 were excluded prior to alignment. Sparse point clouds were generated, cleaned, and optimized using iterative camera calibration and gradual selection to remove low-quality tie points. Dense surface meshes were reconstructed from depth maps using high-quality settings with mild depth filtering, manually cleaned, and textured using mosaic blending. Final models were imported into Autodesk ReCap (Autodesk Inc.) for scaling, orientation, and isolation of individual coral colonies. Mesh artifacts were corrected where necessary, and cleaned models were exported as STL files for subsequent morphometric analyses.

### Morphometric analysis

Morphological metrics were extracted from the 3D coral models, and the following parameters were quantified: substrate coverage (a metric used to quantify horizontal expansion and directly test the SLIPS effect), maximum diameter (defined as the longest horizontal distance across the colony), maximum height (the maximum vertical distance from the base to the highest point), surface area, and total volume. These metrics are consistent with recommended practices for quantifying coral growth using 3D reconstructions (Lange and Perry, 2020; Ferrari et al., 2021). Lastly, percent growth for 6- and 12-month time points were calculated for each metric relative to initial size.

### Micro-computed tomography imaging

High-resolution micro-computed tomography (μCT) was used to quantify fine-scale skeletal development of *Stylophora pistillata* colonies grown on Coral Guard Plugs and conventional calcium carbonate plugs under controlled *ex situ* conditions. At the conclusion of the experiment, coral tissues were removed by immersion in bleach solution for 48 h to isolate the skeletons for imaging. Scans were acquired using a Zeiss Xradia VersaXRM-510 (Carl Zeiss Microscopy GmbH) 3D X-ray microscope at the National Center for Microscopy and Imaging Research, University of California San Diego. Skeletons were scanned through a 360° rotation at an isotropic voxel size of 15 μm (112 kV, 110 μA, 2 s exposure). Three-dimensional reconstructions were generated in Dragonfly (Object Research Systems, Canada), and skeletal density and vertical skeletal extension (measured across the coenosteum) were quantified from the reconstructed volumes. Measurements were compared among Coral Guard Plugs, conventional control plugs, and split-choice Coral Guard Plugs.

### Reflectance measurements

Surface reflectance was measured to quantify optical differences between Coral Guard Plugs and uncoated control plugs following algal fouling. Substrates were submerged in a flow-through aquarium for six weeks to allow natural biofilm development before measurements were performed under water. Reflectance spectra were collected in a black acrylic chamber filled with seawater under homogeneous diffuse illumination provided by a full-spectrum light source (SunWave™ Series, CRI 98, Yuji Lighting). Measurements were acquired using a flat-cut fiber-optic reflectance probe (Ocean Insight; diameter = 0.23 cm) coupled to a miniature spectrometer (Flame, Ocean Insight; boxcar width = 2 nm, spectral resolution = 0.2 nm) following established protocols (Enríquez et al., 2005; Kramer et al., 2023). The probe was positioned 5 mm from the substrate surface at a 45° angle. Five randomly selected surface locations were measured per substrate, and reflectance spectra were normalized to a 99% diffuse reflectance standard (Spectralon, Labsphere).

### Physiological metrics

To evaluate the effects of Coral Guard substrates on coral physiology, symbiont density (cells cm⁻²), total chlorophyll (*a* + *c*₂; µg cm⁻²), soluble protein content (mg cm⁻²), and skeletal density (g cm⁻³) were quantified. Following collection, coral samples were immediately placed in sterile Whirl-Pak bags, frozen at −20 °C, and transported to the University of Hawaiʻi at Mānoa for analysis. Coral tissue was removed from the skeleton using an airbrush with distilled water, homogenized, and subsampled for physiological analyses. Physiological measurements followed established protocols (Wall et al., 2021). Symbiont density was determined using repeated haemocytometer counts (4–8 technical counts per sample). Total chlorophyll (*a* + *c*₂) was extracted from isolated symbiont pellets in 100% acetone at −20 °C for 24 h in darkness and quantified spectrophotometrically using the equations of Jeffrey and Humphrey (1975). Soluble protein content was determined using a bicinchoninic acid (BCA) protein assay (Thermo Fisher Scientific) with bovine serum albumin standards. All physiological metrics were normalized to coral tissue surface area determined using the aluminum foil method (Marsh, 1970). Skeletal density was quantified from the newly deposited encrusting skeleton as described below.

### Statistical analysis

All statistical analyses were performed in R version 4.5.2 (R Core Team, 2025). Growth responses were analyzed separately for each coral species and morphometric parameter. Treatment effects over time were assessed using mixed-effects permutational analysis (MEPA; 1,000 permutations), with treatment and time specified as fixed effects and individual coral fragments included as a random effect to account for repeated measurements. Residual diagnostics were inspected to assess model fit. When significant effects were detected, pairwise permutational *t*-tests were performed. Physiological metrics were analyzed using two-way linear models with species (three levels) and treatment (two levels) as fixed factors. Model assumptions were evaluated by visual inspection of residual and quantile–quantile plots. Type III sums of squares were calculated using the **car** package (Fox & Weisberg, 2019).

## Results

### Effects of Coral Guard on the *ex situ* growth and skeletal morphology of *Stylophora pistillata*

To evaluate the effects of Coral Guard Plugs on coral growth under *ex situ* nursery conditions, *Stylophora pistillata* fragments were grown on Coral Guard Plugs, uncoated control plugs, and split-choice Coral Guard Plugs that enabled simultaneous growth on coated and uncoated surfaces within the same colony (Fig. 2). Coral Guard Plugs significantly enhanced encrusting tissue growth by about 1.7-fold, with substrate coverage reaching 493.93 ± 70.80% compared to 290.11 ± 25.14% on control substrates (permutational t-test, *p* < 0.01; Fig. 2A). Colony diameter was similarly increased, with a 45% significantly higher growth on Coral Guard plugs relative to controls (permutational t-test, *p* < 0.05). Apart from these, other morphometric parameters were unaffected by the Coral Guard.

**Figure 2.**
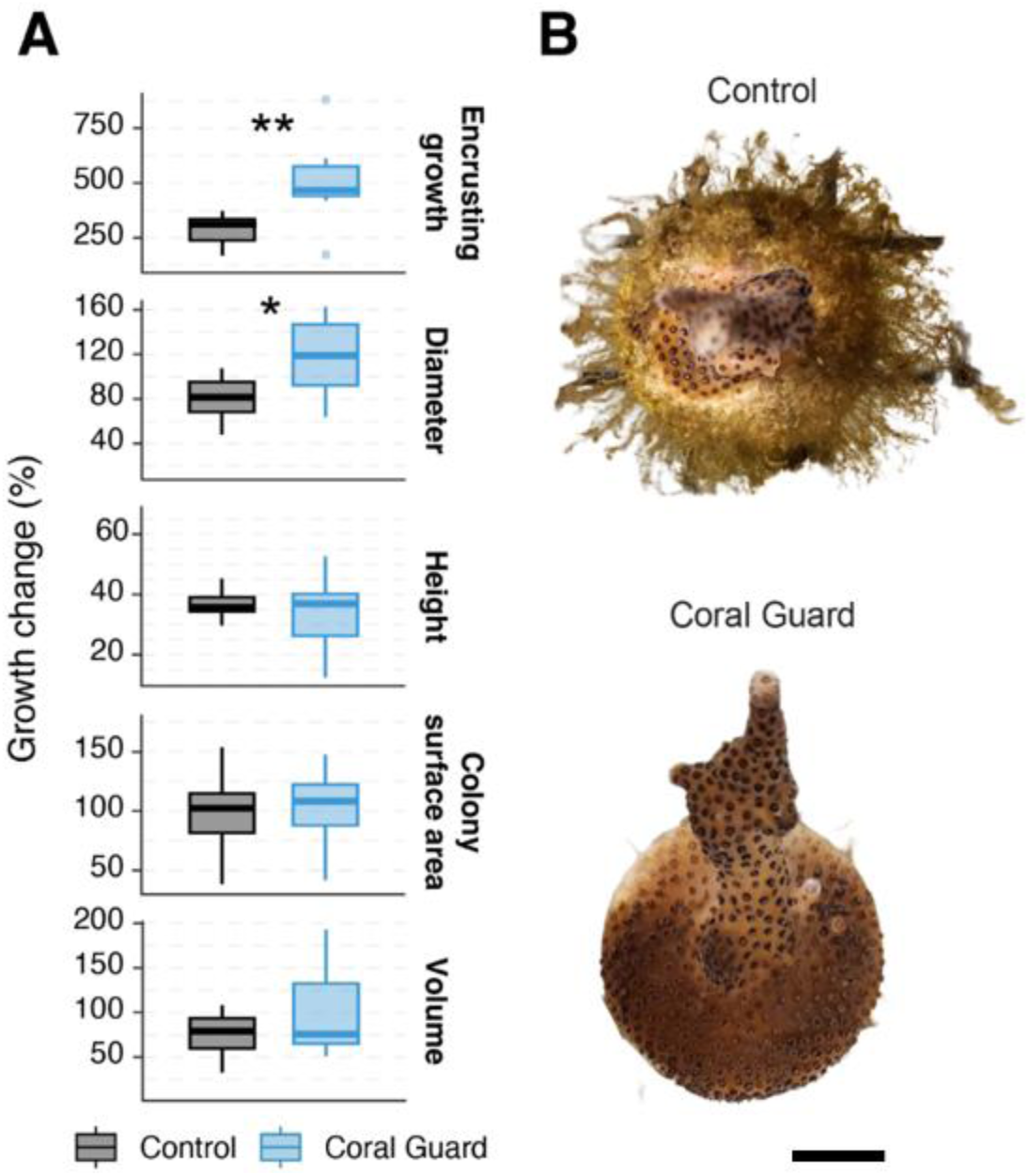
Coral Guard Plugs enhance ex situ growth of *Stylophora pistillata*. (A) Relative growth (%) of *S. pistillata* colonies after 3 months on Coral Guard Plugs and uncoated control plugs. Growth was quantified across five morphometric parameters: substrate coverage, colony diameter, colony height, tissue surface area, and colony volume. Boxplots show the median (horizontal line), interquartile range (box), and whiskers extending to 1.5 × the interquartile range; points represent individual colonies. Asterisks indicate significant differences between treatments (permutational *t*-tests; *P* < 0.05, P < 0.01). (B) Representative photographs of *S. pistillata* colonies grown on uncoated control plugs and Coral Guard Plugs after 3 months (scale bar = 1 cm).

Coral Guard Plugs also altered skeletal development (Fig. 3). μCT analyses revealed a 13% increase in skeletal density relative to controls (1.82 vs. 1.65 g cm⁻³; Fig. 3A) together with a 22% increase in vertical skeletal extension (0.49 ± 0.01 mm versus 0.40 ± 0.01 mm; mean ± SE; Fig. 3B). To determine whether these responses reflected local substrate effects, we evaluated split- choice Coral Guard Plugs, in which individual colonies simultaneously overgrew coated and uncoated surfaces. Skeletal density did not differ between substrate types within the same colony (2% increase on the Coral Guard surface; permutational *t*-test, *p* = 0.94; Fig. 3A). In contrast, vertical skeletal extension remained 17% greater on Coral Guard surfaces than on adjacent control surfaces (0.49 ± 0.02 mm versus 0.42 ± 0.02 mm; Fig. 3B), indicating that Coral Guards locally stimulate skeletal accretion even within individual colonies.

**Figure 3.**
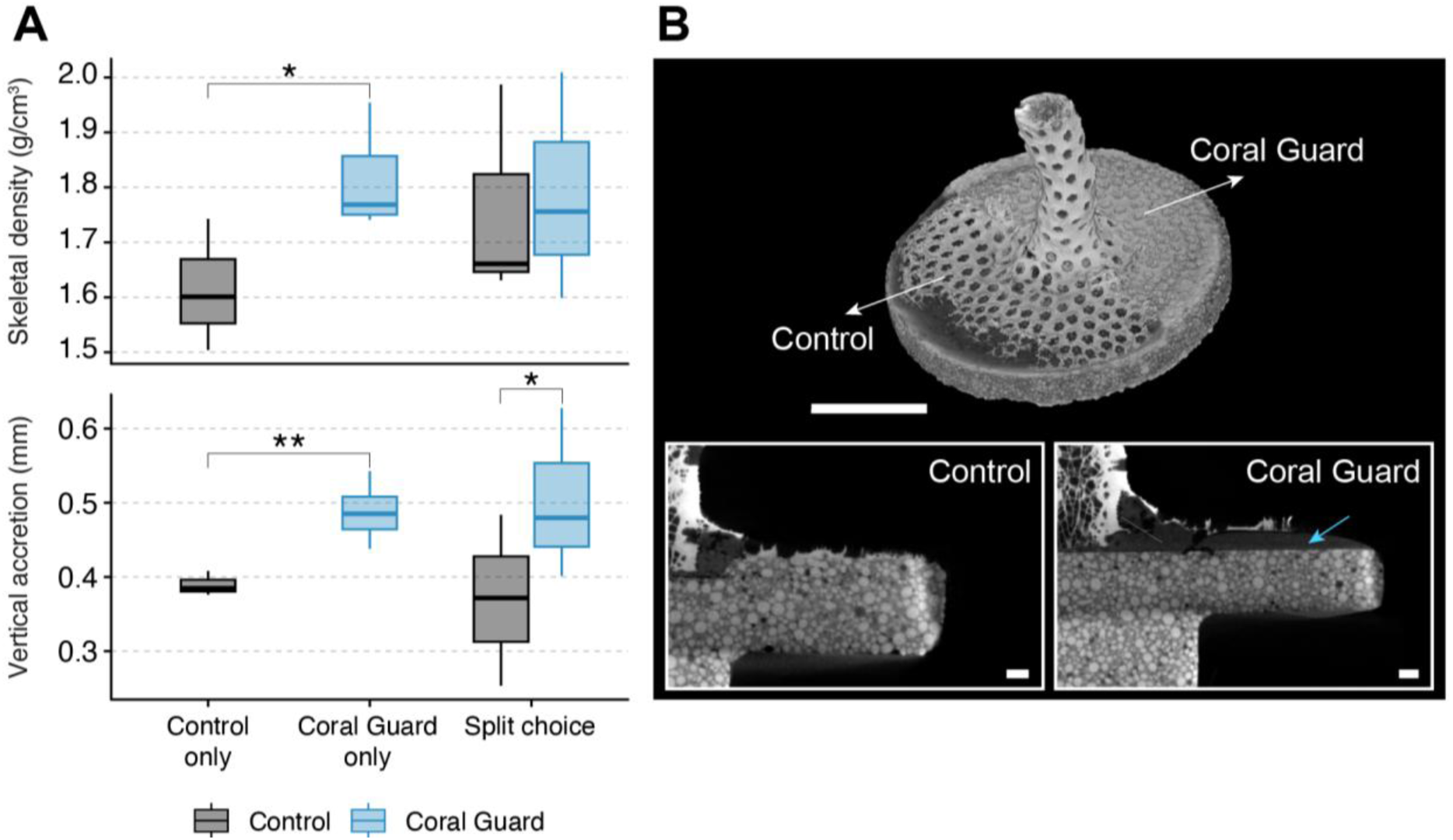
Coral Guard Plugs enhance skeletal development of *Stylophora pistillata* under *ex situ* conditions. (A) Skeletal density (g cm⁻³; top) and vertical skeletal extension (mm; bottom) quantified from μCT reconstructions of colonies grown on uncoated control plugs, split-choice Coral Guard Plugs, and Coral Guard Plugs. Boxplots show the median (horizontal line), interquartile range (box), and whiskers extending to 1.5 × the interquartile range; points represent individual colonies. Asterisks indicate significant differences between treatments (*P* < 0.05, P < 0.01). (B) Representative 3-D μCT reconstruction of a colony grown on a split-choice Coral Guard Plug, illustrating skeletal development across adjacent control and Coral Guard surfaces (top; scale bar = 1 cm). Cross-sectional μCT images highlight differences in vertical skeletal extension between substrate types (bottom; scale bars = 1 mm). The blue arrow indicates the Coral Guard coating layer.

### Reflectivity of Coral Guard substrates

Substrate spectral reflectance (% of incident irradiance) was up to 6-fold higher on Coral Guard substrates (77.8% vs 12.7% in control; 675nm; p < 0.001, permutational t-test) vs control substrates after six weeks of incubation with natural seawater (Fig. 4).

**Figure 4.**
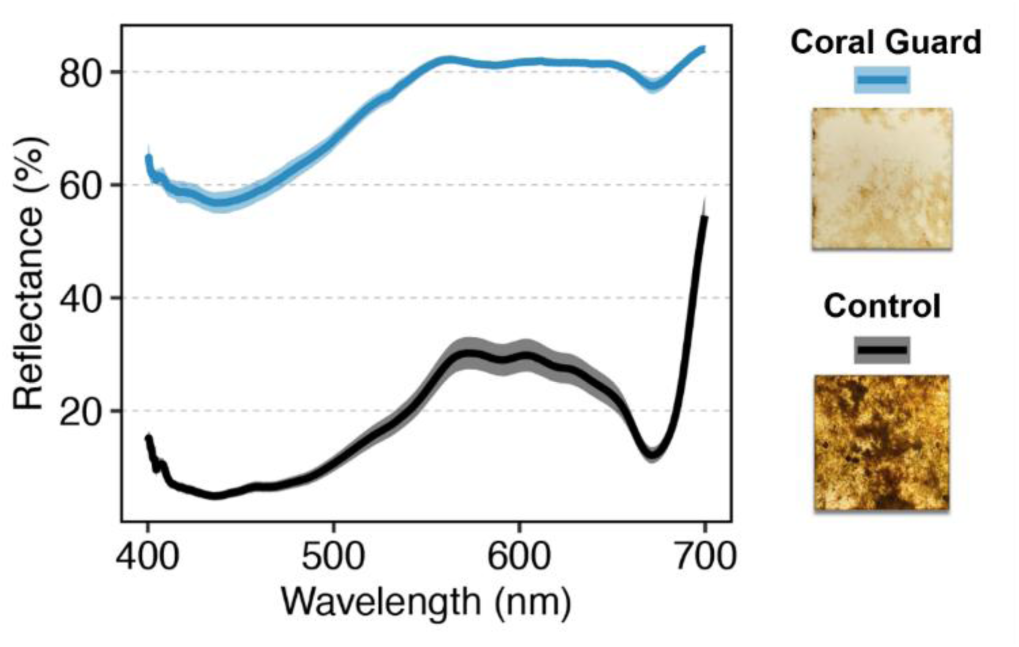
Spectral diffuse reflectance (%) of fouled Coral Guard (blue) and control (black) substrates after six weeks of immersion in a flow-through seawater system. Reflectance was measured across the visible spectrum (400-700 nm). Data represent means ± SE.

### Effects of Fusion Guard Tiles on the *in situ* growth of *Porites evermanni*

Throughout the 12-month deployment, **Fusion Guard Tiles** remained largely free of algal fouling, whereas uncoated control tiles became completely overgrown by turf algae within the first three months (Fig. 5, Fig. S1). This reduction in algal competition was accompanied by significantly enhanced coral growth across multiple morphometric parameters (Fig. 5; permutational ANOVA, *p* < 0.05). After six months, *P. evermanni* colonies on Fusion Guard Tiles exhibited 2.6-fold greater lateral tissue expansion than colonies on control tiles (194.76 ± 31.03% versus 74.42 ± 12.06%; permutational *t*-test, *P* < 0.01; Fig. 5A). Likewise, three-dimensional tissue surface area increased by more than 2.7-fold (87.23 ± 16.17% versus 32.08 ± 3.12%; permutational *t*-test, *P* < 0.01). Consistent with these enhanced growth rates, microfragment fusion was already evident on Fusion Guard Tiles after six months, whereas no fusion occurred on control tiles (Fig. 5B; Fig. S2).

**Figure 5.**
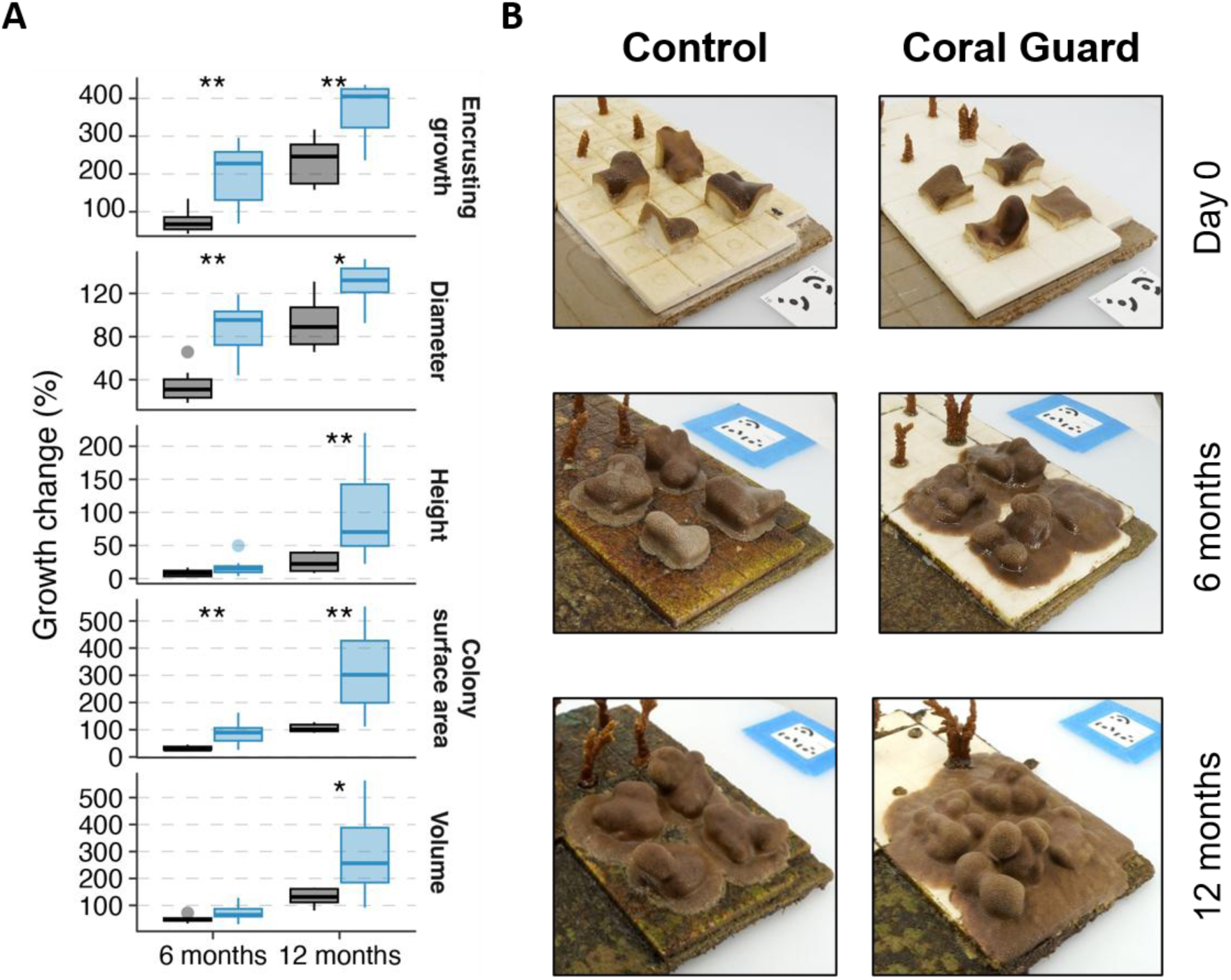
(A) Growth rates of *Porites evermanni* fragments on Control and Coral Guard tiles at the 6- and 12-month time points. Boxplots depict the median (*horizontal line*), interquartile range (first and third quartiles), and whiskers as ±1.5 interquartile range with dots representing outliers. Asterisks denote significant differences between treatments within specific time points and metrics based on permutational t-tests (*\*p* < 0.05, \*\**p* < 0.01, \*\*\**p* < 0.001). **(B)** Representative images of the *P. evermanni* plates at the starting, 6-month and 12-month time point.

After 12 months, colonies on Fusion Guard Tiles continued to exhibit significantly greater growth across multiple morphometric parameters than colonies on control tiles (Fig. 5). Lateral tissue expansion remained approximately 1.6-fold greater on Fusion Guard Tiles (367.62 ± 27.82% versus 232.33 ± 23.90%; permutational *t*-test, *P* < 0.01), while 3D tissue surface area increased by approximately 3.0-fold (317.02 ± 60.77% versus 106.08 ± 5.82%; permutational *t*-test, *P* < 0.01). Vertical extension was approximately 4-fold greater on Fusion Guard Tiles than on control tiles (99.49 ± 28.85% versus 24.82 ± 5.60%; permutational *t*-test, *P* < 0.01), and colony volume was approximately 2.2-fold greater (*P* < 0.05). By 12 months, microfragments on Fusion Guard Tiles had fused into continuous colonies, whereas fusion on control tiles had only begun (Fig. 5B).

### Effects of Fusion Guard Tiles on the physiology and skeletal density of *Porites evermanni*

After 12 months, *Porites evermanni* colonies grown on Fusion Guard Tiles exhibited significantly greater symbiont densities than colonies grown on control tiles, with an approximately 1.7-fold increase (*P* < 0.05, permutational ANOVA; Fig. 6). Soluble protein content was also significantly higher, increasing by approximately 2.5-fold on Fusion Guard Tiles relative to controls (*P* < 0.001). In contrast, total chlorophyll (*a* + *c*₂) content did not differ significantly between treatments, although greater variability was observed among control colonies. Skeletal density of newly deposited skeleton was approximately 10% higher in colonies grown on Fusion Guard Tiles than in controls (*P* < 0.05).

**Figure 6.**
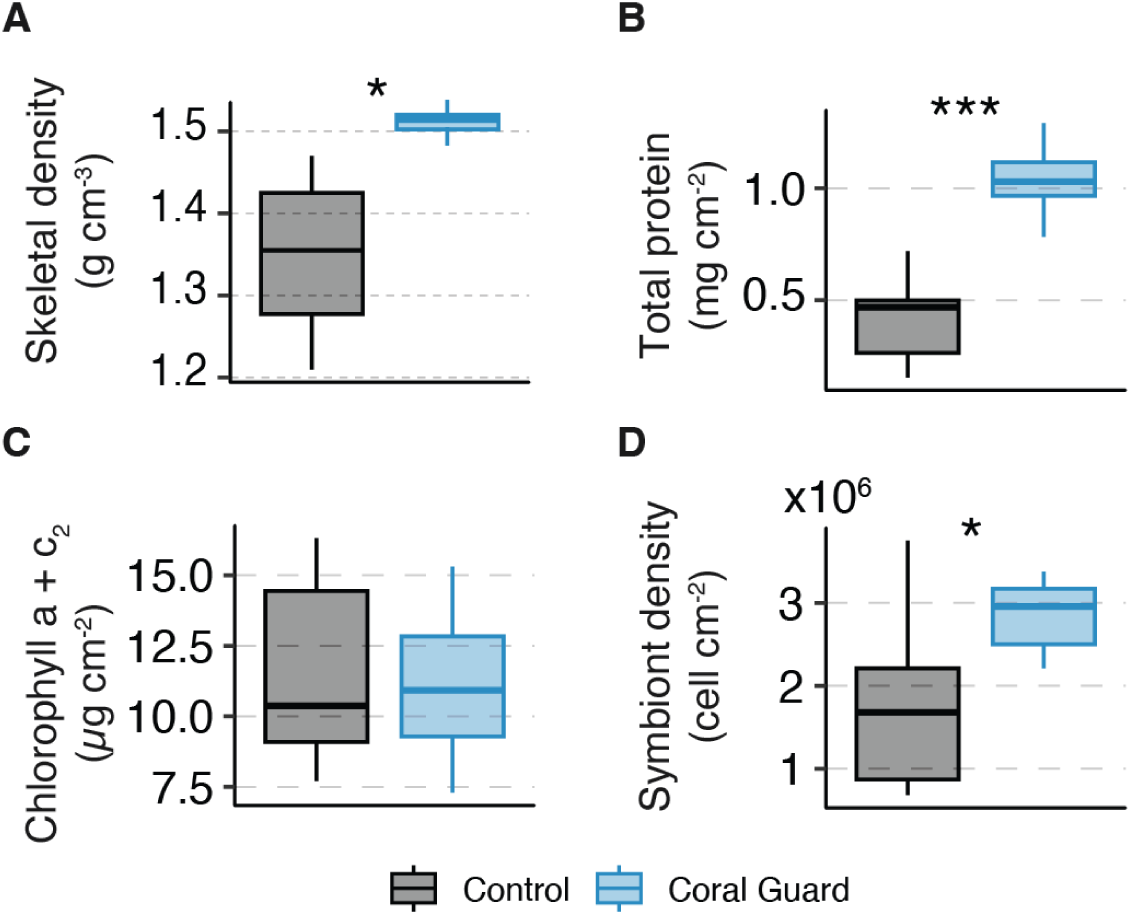
Fusion Guard Tiles enhance the physiological condition and skeletal density of *Porites evermanni* after 12 months of *in situ* nursery growth. (A) Symbiont density (cells cm⁻²), (B) soluble protein content (mg cm⁻²), (C) total chlorophyll (*a* + *c*₂) (µg cm⁻²), and (D) skeletal density (g cm⁻³) of *P. evermanni* colonies grown on Fusion Guard Tiles and uncoated control tiles. Boxplots show the median (horizontal line), interquartile range (box), and whiskers extending to 1.5 × the interquartile range; points represent individual colonies. Asterisks indicate significant differences between treatments (permutational *t*-tests; *P* < 0.05, \*\**P* < 0.001).

## Discussion

Coral reef restoration is increasingly recognized as a critical intervention for sustaining reef structure and ecosystem function under accelerating climate change and local environmental stressors. However, the large-scale propagation of asexually derived coral outplants remains constrained by major operational bottlenecks, including high maintenance requirements, intensive manual labor, slow growth, and often low post fragmentation survivorship. These limitations are strongly influenced by environmental conditions (Foo et al, 2021), species specific responses (Lustic et al, 2020; Henley et al, 2022), and local ecological context (Pausch et al, 2018), but across many restoration systems, frequent cleaning and management of algal fouling remain among the most labor intensive and costly aspects of nursery operations. Overcoming these bottlenecks will require technologies that not only enhance coral growth and survivorship but also reduce maintenance demands and increase restoration throughput.

In this study, we demonstrate that the Coral Guard platform fundamentally modifies the coral–substrate interface, resulting in sustained suppression of algal fouling, accelerated tissue expansion, and enhanced microfragment fusion under *in situ* coral nursery conditions (Figs. 1, 5, Fig. S1). Across complementary *ex situ* and *in situ* experiments, these effects translated into substantial improvements in coral growth, highlighting substrate microenvironment engineering as a scalable strategy to improve restoration efficiency. The most consistent response observed across experiments was the suppression of benthic algal fouling on Coral Guard surfaces. In the *in situ* nursery, Fusion Guard Tiles remained largely free of algal overgrowth throughout the 12- month deployment, whereas uncoated control tiles were rapidly colonized by turf algae within the first three months (Fig. 5, Fig. S1). Benthic algae are well known to suppress coral growth by competing for space and light and by altering local biogeochemical conditions through the release of dissolved organic carbon and stimulation of microbial activity (Haas et al. 2011; Haas et al. 2016; Barott & Rohwer 2012). By preventing algal establishment, Coral Guard surfaces effectively remove this competitive barrier, creating a substrate microenvironment that favors coral tissue expansion and colony development.

Using 3D photogrammetry, we observed substantial positive effects of the Coral Guard platform on coral growth and colony development. In *Porites evermanni*, Fusion Guard Tiles promoted rapid lateral tissue expansion, with substrate coverage increasing 2.6-fold relative to control tiles within the first 6 months, followed by sustained increases across all measured morphometric parameters after 12 months, including three-dimensional (3D) tissue surface area, colony volume, and vertical extension (Fig. 5). Complete microfragment fusion was achieved within 12 months on Fusion Guard Tiles, whereas fusion on control tiles remained delayed and incomplete. Accelerated microfragment fusion is a particularly important outcome for coral restoration because it shortens nursery residence times and promotes the formation of larger, structurally stable colonies (Page et al. 2018; Knapp et al. 2022). Earlier colony fusion may also facilitate earlier reproductive maturity, which in many coral species is closely linked to colony size (Okubo et al. 2007).

An important finding of this study was that enhanced coral growth on the Coral Guard platform was not associated with reductions in skeletal density. In contrast to the commonly observed trade-off between rapid coral growth and reduced skeletal density (Lough & Barnes, 2000; Carricart-Ganivet, 2004), both *Porites evermanni* under *in situ* nursery conditions and *Stylophora pistillata* under controlled *ex situ* conditions exhibited simultaneous increases in skeletal growth and skeletal density. In *P. evermanni*, colonies grown on Fusion Guard Tiles exhibited approximately 10% higher skeletal density than colonies grown on control tiles after 12 months (Fig. 6). Similarly, μCT analyses of *S. pistillata* revealed approximately 13% higher skeletal density together with significantly greater vertical skeletal extension on Coral Guard Plugs than on control plugs (Fig. 3). Together, these findings indicate that the Coral Guard platform enhances skeletal development without compromising skeletal density.

One potential concern was that enhanced skeletal development on the Coral Guard platform might occur at the expense of tissue condition or symbiont performance through energetic trade-offs (Anthony et al., 2002; Spalding et al., 2017; Wall et al., 2018). However, physiological analyses did not indicate any adverse effects associated with Coral Guard surfaces (Fig. 6A–D). Instead, *Porites evermanni* colonies grown on Fusion Guard Tiles exhibited approximately 1.7- fold greater symbiont densities and 2.5-fold higher host tissue protein content than colonies grown on control tiles after 12 months (Fig. 6). Elevated protein content likely reflects greater host tissue biomass and improved colony condition. Despite significantly higher symbiont densities, total chlorophyll concentrations normalized to tissue surface area remained similar between treatments. This pattern is consistent with photoacclimation under increased local light availability (Kinzie et al., 1984; Kuguru et al., 2010).

Several interacting mechanisms may contribute to these responses. By suppressing turf algal overgrowth, the Coral Guard platform likely reduces competitive stress and the associated microbial and biogeochemical disturbances at the coral–substrate interface, thereby lowering the energetic costs of benthic competition (Barott & Rohwer, 2012; Chadwick & Morrow, 2011; Haas et al., 2011, 2016). In addition, Coral Guard surfaces maintained substantially higher reflectance than algal-fouled control surfaces (Fig. 4), increasing diffusely scattered light available for photosynthesis (Wangpraseurt et al., 2014). Previous studies have shown that substrate reflectance can substantially influence the benthic light environment experienced by corals *in situ* (Enríquez et al., 2005; Wangpraseurt et al., 2014). Together, reduced benthic competition and an enhanced local light microenvironment may allow corals to simultaneously support tissue growth, skeletal development, and colony expansion without apparent physiological trade-offs.

It is important to recognize that coral growth form and competitive interactions are highly species-specific, and consequently the benefits of the Coral Guard platform are likely to vary among coral taxa and morphologies. Corals with strongly encrusting growth forms, or branching species that initially prioritize lateral tissue expansion before vertical growth, may benefit most from the creation of competition-free substrate space. Although preliminary, additional experiments with *Montipora capitata* and *Pavona varians* revealed species-specific responses. *M. capitata* showed qualitatively positive responses, including greater host tissue protein content, whereas *P. varians* exhibited no detectable treatment effects. These observations suggest that benthic algal competition was not the primary factor limiting growth in *P. varians* under our experimental conditions, and that reducing competition at the coral–substrate interface therefore had less influence on colony development. More broadly, coral growth reflects the net outcome of multiple interacting environmental drivers, including light availability, water flow, heterotrophy, sedimentation, and benthic competition. Consequently, the magnitude of Coral Guard benefits will likely depend on coral species, colony morphology, and environmental context.

From a practical coral restoration perspective, these findings address a central operational bottleneck: the need for frequent manual cleaning to control algal fouling in coral nurseries. Coral gardening and microfragmentation approaches remain constrained by high labor demands, maintenance costs, and limited biological throughput, restricting restoration scalability (Boström- Einarsson et al. 2020; Schmidt-Roach et al. 2025). By passively suppressing algal overgrowth, Coral Guards reduce maintenance requirements while simultaneously enhancing coral growth, tissue expansion, and microfragment fusion. This combination has the potential to shorten nursery residence times, accelerate colony formation, and increase restoration throughput. More broadly, our findings demonstrate that engineering the coral–substrate microenvironment represents a powerful and underutilized strategy for coral restoration. Rather than directly manipulating coral physiology, Coral Guards modify the competitive and optical environment at the benthic interface in favor of coral growth and development, highlighting substrate microenvironment engineering as a scalable, low-maintenance approach for advancing coral nursery production and reef restoration.

## Acknowledgements

This research was supported by the Defense Advanced Research Projects Agency (DARPA) under the Reefense Program (award HR0011-22-C-0134; D.W., R.T.), the Gordon and Betty Moore Foundation through the Moore Inventor Fellowship (#13624, D.W.), the Accelerating Innovation to Market Grant at the University of California San Diego (D.W.), and the CA CARES Grant (D.W.), a Climate Impact initiative of the University of California. We thank Ben Jones and the members of the R3D Consortium for logistical and administrative support throughout the project. The views, opinions, and findings expressed in this article are those of the authors and should not be interpreted as representing the official views or policies of the U.S. Department of Defense or the U.S. Government.

## Disclosure statement

While no competing interests exist for any of the authors, the following information is provided for the sake of full transparency: D.W., S.Ko. are authors on a provisional patent application related to this work. D.W. is founder of Hybrid Reef Solutions and is an advisory board member of the Coral Restoration Consortium, community practice for reef restoration. The authors declare that they have no other competing interests.

## Data and Code Availability

All raw data presented in this paper is freely available (Figshare link: xxx)

## Author Contributions

**Conceived the study:** S.Ko., C.S., R.T., and D.W. conceived the *in situ* coral nursery study, while S.Ko., S.Ku., and D.W. conceived the *Stylophora* growth study and coral choice experiment. **Performed the study:** S.Ko. designed and fabricated the Coral Guard substrates and led the *in situ* coral nursery and *ex situ* growth experiments, with contributions from C.S., J.K., A.P., L.B., N.L., S.Ku., A.B., E.S., C.W., and S.M. **Analyzed and interpreted the data:** N.K. and S.Ko. analyzed the data and generated the figures. N.K., S.Ko., and D.W. interpreted the data. **Acquired funding and supervised the study:** D.W. **Wrote the manuscript:** S.Ko., N.K., and D.W. wrote the manuscript with input from all authors. All authors reviewed and approved the final manuscript.

