## Supplementay information for "Coral Guard substrates accelerate growth and fragment fusion for coral restoration"

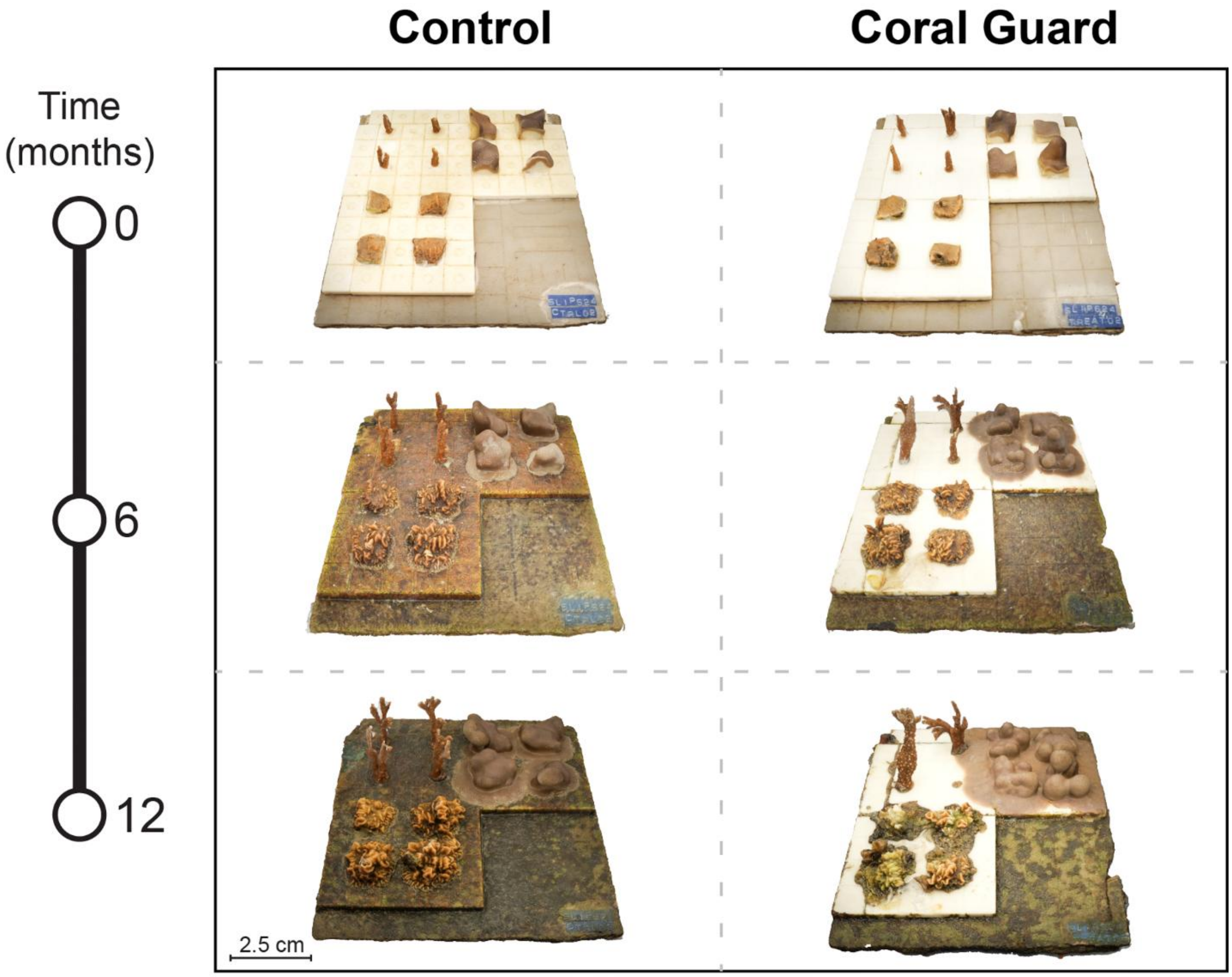


**Fig. S1. Representative photogrammetric reconstructions showing algal fouling progression on Control and Fusion Guard Tiles over a 12-month *in situ* coral nursery deployment.** Representative experimental tiles are shown at 0, 6, and 12 months for the Control and Fusion Guard treatments.

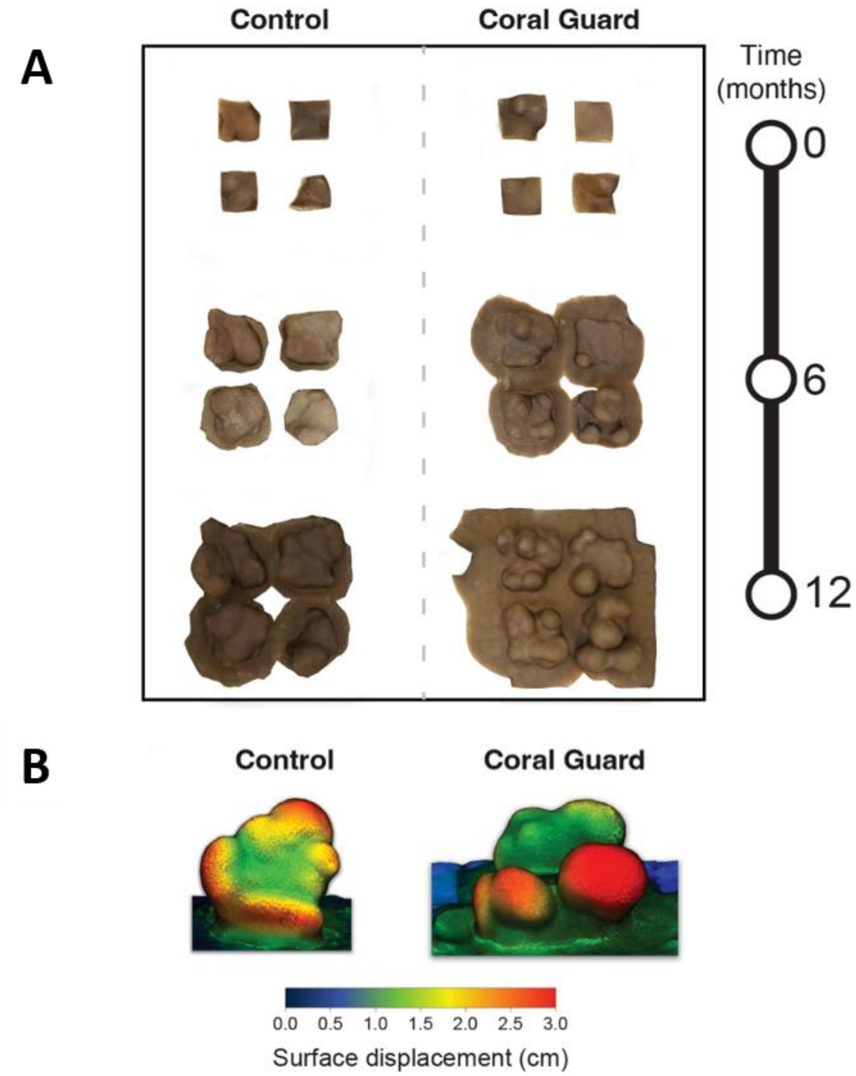


**Fig. S2. Visualization of *Porites evermanni* growth and microfragment fusion on Control and Fusion Guard Tiles.** (A) Representative top-down views of coral tissue after segmentation, with the tiles and background removed to highlight tissue expansion and fragment fusion at 0, 6, and 12 months. (B) Representative three-dimensional photogrammetric reconstructions of colonies grown on Control and Fusion Guard Tiles. Surface displacement maps illustrate regions of greatest outward tissue extension relative to the initial colony geometry.

**Supplementary observations for *Montipora capitata* and *Pavona varians***

In addition to *Porites evermanni*, *in situ* coral nursery experiments were conducted using *Montipora capitata* and *Pavona varians* on Fusion Guard Tiles and uncoated control tiles. These datasets were excluded from the main analyses because substantial fragment loss during the experimental period resulted in insufficient replication for robust quantitative comparisons at the 12-month endpoint.

Following deployment, uncoated control surfaces, together with all unprotected regions of the Fusion Guard Tiles, rapidly developed dense turf algal fouling. By 3 months, virtually all unprotected surfaces were completely overgrown, whereas Coral Guard-coated regions remained largely free of fouling and retained their initial appearance and reflectivity throughout the 12-month deployment (Fig. S1). Two localized coating delamination events were observed, including one major delamination on an *M. capitata* tile and one minor delamination on a *P. varians* tile. Despite partial coating detachment, fouling beneath the affected regions remained limited, and all remaining Coral Guard surfaces remained intact and functional throughout the study.

Coral survivorship remained high during the first 6 months of deployment. During this period, only a single *M. capitata* fragment detached from each treatment, while only one *P. evermanni* fragment mortality occurred in the control treatment. Following the 6-month survey, however, the nursery infrastructure and experimental arrays were relocated to a nearby facility. This relocation coincided with substantial fragment loss and stress responses in both *M. capitata* and *P. varians*. Four additional *M. capitata* fragments detached from Fusion Guard Tiles during handling, substantially reducing replication, while four *P. varians* fragments in each treatment bleached following relocation and did not recover before the end of the experiment. In contrast, *P. evermanni* remained comparatively stable, with seven surviving fragments remaining in both treatments after 12 months (Fig. S1).

Consequently, the combined effects of fragment detachment, bleaching-associated mortality, and reduced replication precluded robust quantitative evaluation of treatment effects in *M. capitata* and *P. varians*. Nevertheless, qualitative observations during the first 6 months suggested species-specific differences in response to Coral Guards. *Montipora capitata* exhibited trends broadly consistent with the enhanced growth and reduced fouling observed for *P. evermanni*, whereas *P. varians* showed comparatively limited responses under the conditions tested. These observations suggest that the benefits of Coral Guards may vary among coral species, growth forms, and environmental contexts (Fig. S3).

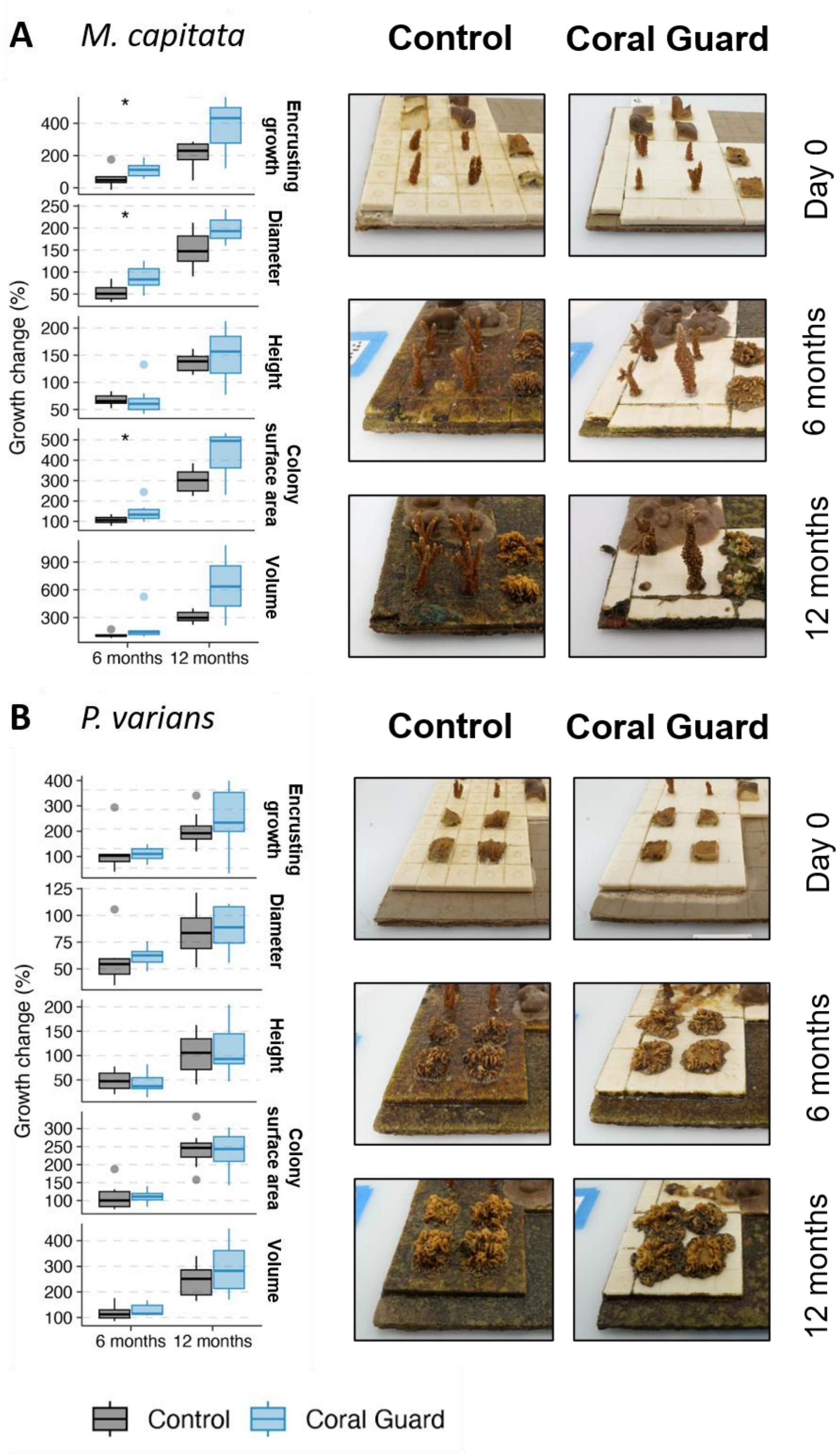

**Figure S3. Supplementary growth responses of *Montipora capitata* (A) and *Pavona varians* (B) grown on Fusion Guard Tiles and uncoated control tiles during a 12-month *in situ* coral nursery experiment.** Relative growth (%) is presented for five morphometric parameters: substrate coverage, colony diameter, colony height, three-dimensional (3D) tissue surface area, and colony volume. Boxplots show growth at the 6- and 12-month sampling points, with representative colony images for each treatment. Boxplots depict the median (horizontal line), interquartile range (box), and whiskers extending to 1.5 × the interquartile range; points represent individual colonies. Asterisks indicate significant differences between treatments (permutational *t*-tests; $P < 0.05$, P < 0.01, **$P < 0.001$). Owing to fragment loss and bleaching following nursery relocation, these data are presented for qualitative comparison only and were not included in the primary statistical analyses.

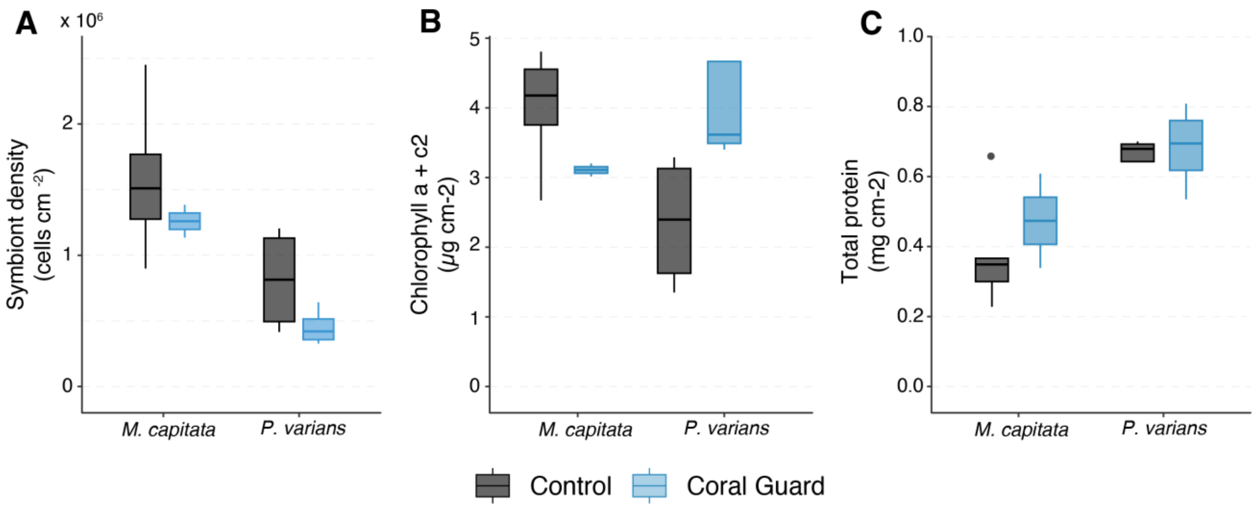


**Figure S4. Supplementary physiological responses of *Montipora capitata* and *Pavona varians* after 12 months of growth on Fusion Guard Tiles and uncoated control tiles.** (A) Symbiont density (cells $cm^{-2}$), (B) total chlorophyll ($a + c_2$) (µg $cm^{-2}$), and (C) soluble protein content (mg $cm^{-2}$). Boxplots show the median (horizontal line), interquartile range (box), and whiskers extending to 1.5 × the interquartile range; points represent individual colonies. Owing to fragment loss and bleaching following nursery relocation, these data are presented for qualitative comparison and were not included in the primary statistical analyses.

**Supplementary Table 1:** List of R3D consortium members. Lab leads are noted in bold.

| Author | Institution | ORCID |
|---|---|---|
| **Benjamin A. Jones** | Applied Research Laboratory at the University of Hawai'i | 0009-0000-2692-7443 |
| Joshua Levy | Applied Research Laboratory at the University of Hawai'i | |
| Sean Mahaffey | Applied Research Laboratory at the University of Hawai'i | |
| Aricia Argyris | Applied Research Laboratory at the University of Hawai'i | |
| Mark Aruda | Applied Research Laboratory at the University of Hawai'i | |
| Ian Robertson | Applied Research Laboratory at the University of Hawai'i | |
| **Zhenhua Huang** | University of Hawai'i at Mānoa | 0000-0001-6665-7230 |
| Ayrton Medina-Rodriguez | University of Hawai'i at Mānoa | 0000-0002-0666-9472 |
| Mert Gokdepe | University of Hawai'i at Mānoa | |
| Brady Halvorson | University of Hawai'i at Mānoa | |
| Jon Chase | University of Hawai'i at Mānoa | |
| Charlotte White | University of Hawai'i at Mānoa | |
| Cami Dillon | University of Hawai'i at Mānoa | |
| Kristian McDonald | University of Hawai'i at Mānoa | |
| Anna Mikkelsen | University of Hawai'i at Mānoa | |
| **Josh Madin** | Hawai'i Institute of Marine Biology | 0000-0002-5005-6227 |
| Mollie Asbury | Hawai'i Institute of Marine Biology | |
| Jessica Reichert | Hawai'i Institute of Marine Biology | 0000-0003-2245-4188 |
| Hendrikje Jorissen | Hawai'i Institute of Marine Biology | |
| Nina Schiettekatte | Hawai'i Institute of Marine Biology | |
| Marion Chapeau | Hawai'i Institute of Marine Biology | |
| **Rob Toonen** | Hawai'i Institute of Marine Biology | 0000-0001-6339-4340 |
| Christopher R. Suchocki | Hawai'i Institute of Marine Biology | 0000-0001-6811-1987 |
| Van Wieringen | Hawai'i Institute of Marine Biology | 0000-0002-6256-4018 |
| Chris Jury | Hawai'i Institute of Marine Biology | |
| Daniel Schar | Hawai'i Institute of Marine Biology | |
| Madeleine Hardt | Hawai'i Institute of Marine Biology | |
| Claire Lewis | Hawai'i Institute of Marine Biology | 0000-0003-1081-2734 |
| Claire Bardin | Hawai'i Institute of Marine Biology | |
| Joshua Kualani | Hawai'i Institute of Marine Biology | |
| **Crawford Drury** | Hawai'i Institute of Marine Biology | 0000-0001-8853-416X |
| **Kira Hughes** | Hawai'i Institute of Marine Biology | |
| Josh Hancock | Hawai'i Institute of Marine Biology | 0000-0002-4814-1385 |
| Carlo Caruso | Hawai'i Institute of Marine Biology | |
| **Andrea Grottoli** | Ohio State University | 0000-0001-6053-9452 |
| Shannon Dixon | Ohio State University | 0009-0007-9882-7936 |
| Ann Marie Hulver | Ohio State University | 0000-0003-0466-9070 |
| **Joshua D. Voss** | Florida Atlantic University | 0000-0002-0653-2767 |
| Allison Klein | Florida Atlantic University | 0009-0004-2670-4222 |
| **Siddhartha Verma** | Florida Atlantic University | 0000-0002-8941-0633 |

| **Author** | **Institution** | **ORCID** |
|---|---|---|
| Alejandro Alvaro | Florida Atlantic University | |
| **Richard Argall** | Makai Ocean Engineering | |
| Kevin Chun | Makai Ocean Engineering | |
| William Hicks | Makai Ocean Engineering | |
| Alex LeBon | Makai Ocean Engineering | |
| John Yeh | Makai Ocean Engineering | |
| **Aaron Thode** | Scripps Institution of Oceanography, UC San Diego | |
| Oceane Boulais | Scripps Institution of Oceanography, UC San Diego | |
| **Daniel Wangprasert** | Scripps Institution of Oceanography, UC San Diego | 0000-0003-4834-8981 |
| Samapti Kundu | Scripps Institution of Oceanography, UC San Diego | |
| Natalie Levy | Scripps Institution of Oceanography, UC San Diego | |
| Lindsey Badder | Scripps Institution of Oceanography, UC San Diego | |
| Stefan Kolle | Scripps Institution of Oceanography, UC San Diego | 0000-0003-2024-9690 |
